## Supplementary tables and figures for "Intrinsically disordered CsoS2 acts as a general molecular thread for α-carboxysome shell assembly"

for

List of Supplementary materials

Supplementary Table 1-3
Supplementary Figure S1 – S11

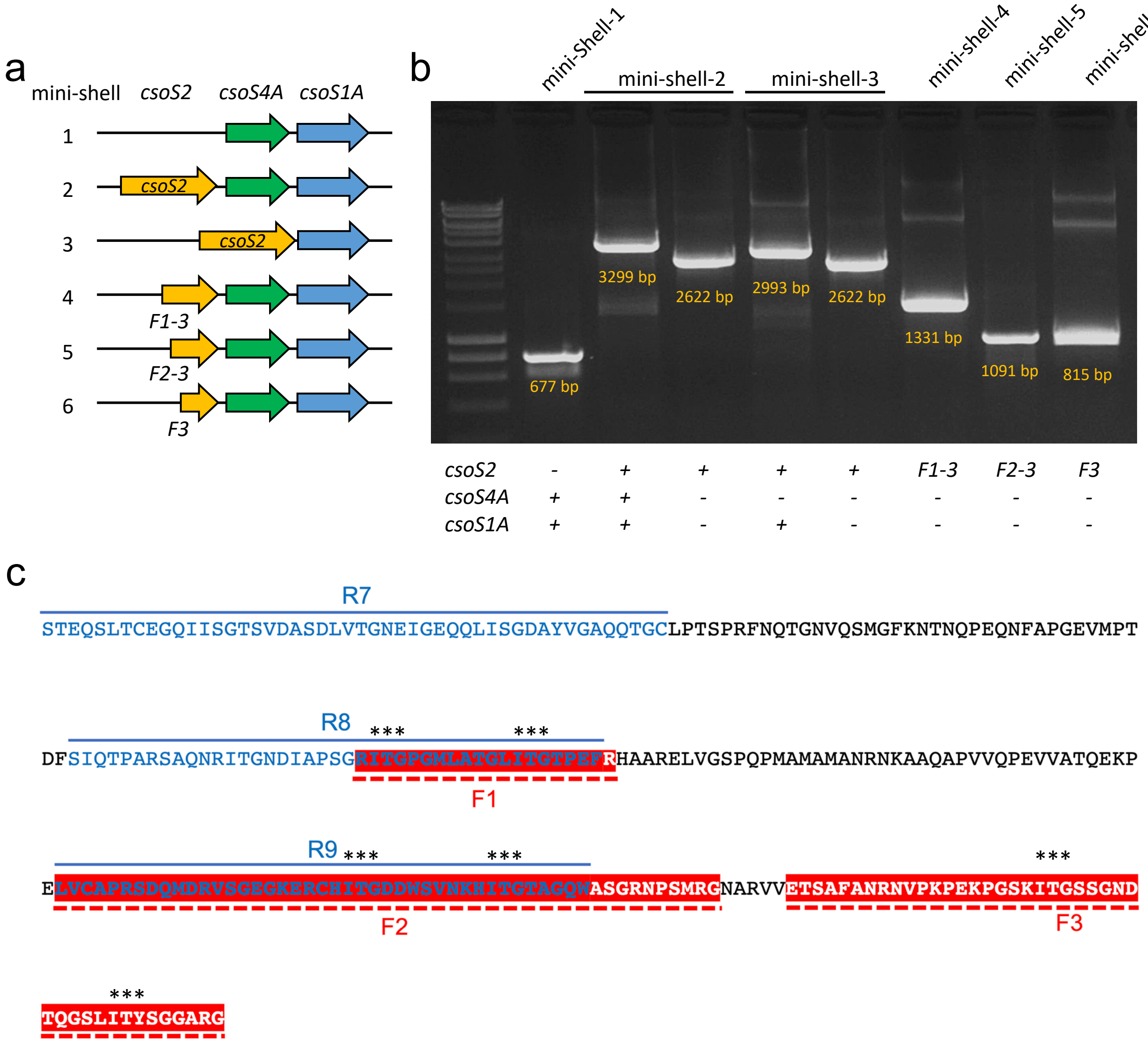


**Figure S1** | Construction of the different mini-shell forms. (a) The genetic arrangements of mini-shell-1 to mini-shell-6 constructs generated in this study. (b) PCR verification of the mini-shell constructs using the primers listed in Supplementary Table 3. The genes in PCR products are indicated at the bottom of the gel image. Two sets of primers are used in the mini-shell 2 and 3 constructs. The sizes (bp) of PCR products are labelled in orange. (c) The protein sequence of CsoS2 C-terminal domain. The three interaction fragments in the C-terminal region (F1, F2, F3) newly identified in the $T$ = 9 shell are shown in red. The three additional repeats in the C-terminal region (R7, R8, R9) previously identified (1) are represented in blue. *** indicates the I(V)TG motif, which was replaced by AAA in the CsoS2-Cm mutant.

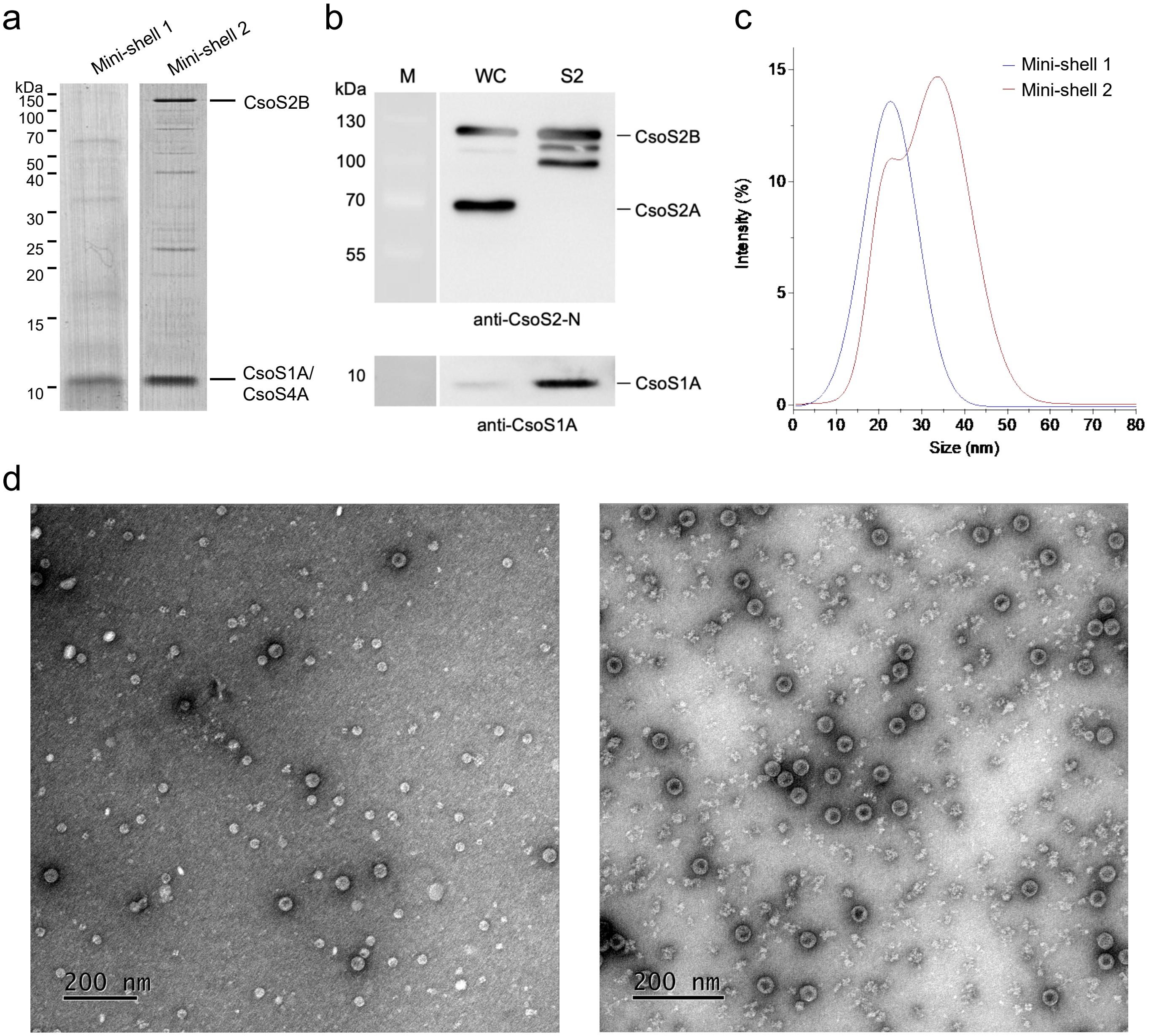


**Figure S2** | Characteristics of shells generated from mini-shell 1 (CsoS4A-CsoS1A) and mini-shell 2 (CsoS2-CsoS4A-CsoS1A) constructs. (a) SDS-PAGE results revealed the major protein components of purified shells from mini-shell-1 and mini-shell-2. (b) Immunoblot analysis using anti-CsoS2 antibody (GenScript, USA) revealed that both CsoS2A and CsoS2B were expressed in the *E. coli* mini-shell 2 construct, but only CsoS2B was incorporated into the mini-shells and CsoS2A was not detectable. WC: whole cell lysate of the CsoS2-CsoS4A-CsoS1A mini-shell 2 construct; S2: isolated CsoS2-CsoS4A-CsoS1A mini-shells. (c) Dynamic light scattering (DLS) analysis of shell sizes from mini-shell 1 and mini-shell 2. (d) Electron microscopy (EM) images of negatively stained purified shells from mini-shell 1 and mini-shell 2.

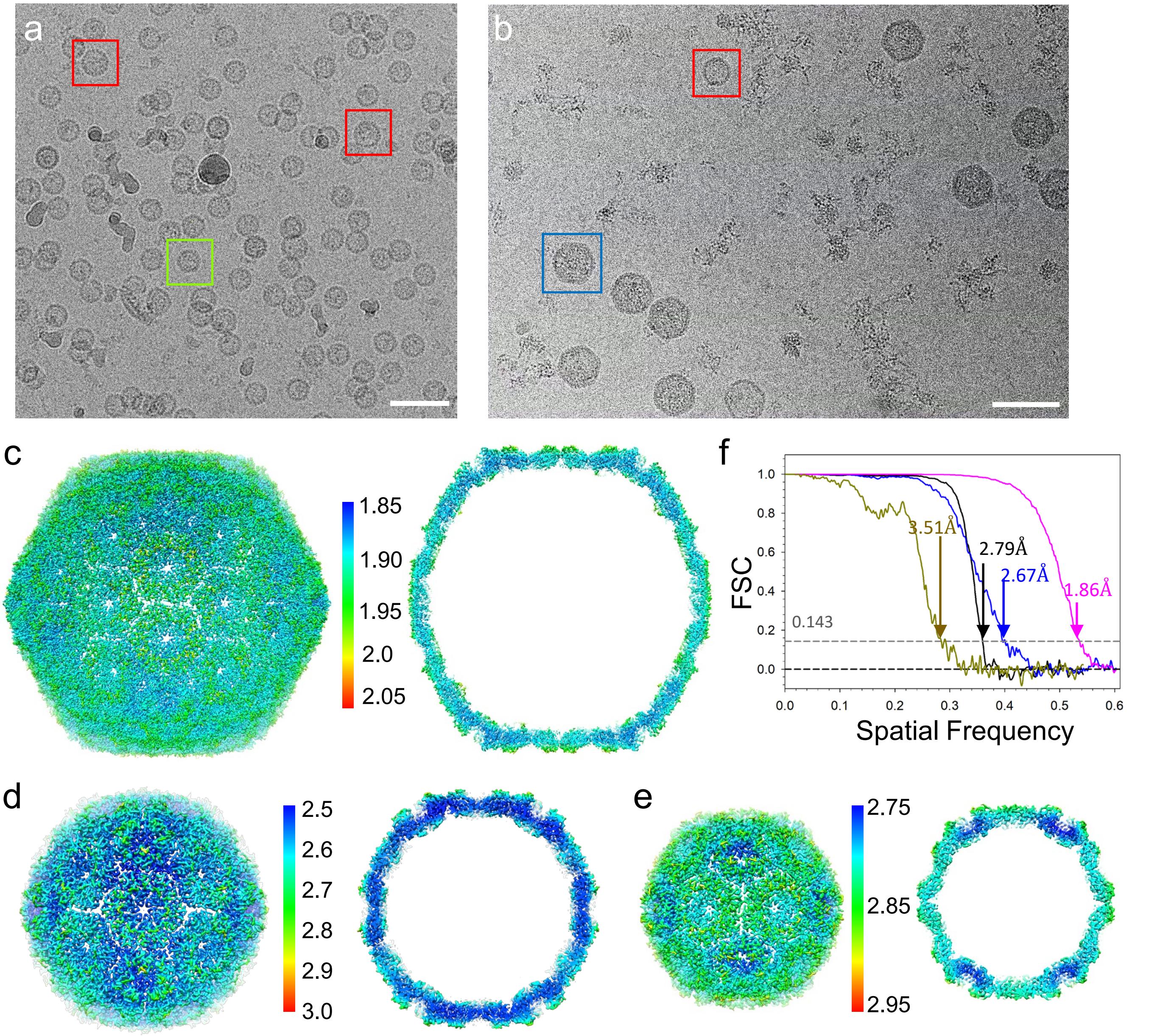


**Figure S3** | CryoEM data processing. (a-b) Representative micrographs of shells produced from mini-shell 1 (a) and mini-shell 2 (b), respectively. Boxed particles have different sizes: blue, large shells (*T* = 9); red, medium shell (*T* = 4); and green, small shell (*T* = 3). Scale bars: 50 nm. (c-e) CryoEM maps of *T* = 9 (c), *T* = 4 (d) and *T* = 3 (e), shown in top view (left) and central slice (right). Maps are coloured according to their local resolutions. (f) Fourier Shell Correlation (FSC) of shells, *T* = 9 in magenta, *T* = 4 in blue (from mini-shell 2) and brown (from mini-shell 1), and *T* = 3 in black, with resolutions indicated at FSC=0.143 cut-off.

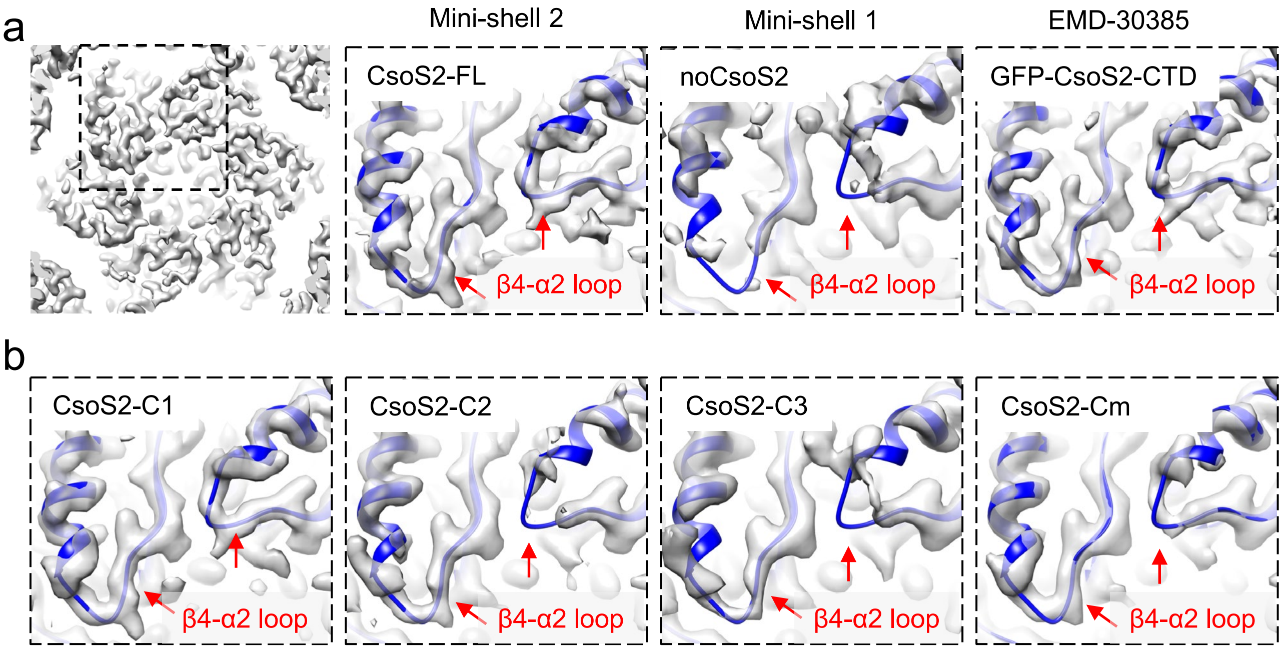


**Figure S4** | Comparison of $T$ = 4 shells with and without CsoS2 from different mini-shell constructs. (a) Comparison of $T$ = 4 shell hexamer maps (gray) from mini-shell-1, mini-shell-2, and a GFP-CsoS2-CTD/4A-1A (EMD-30385). The major difference in hexamer density is located in the loop between β4 and α2 in S1A (red arrows), which could not be resolved in the shell without CsoS2. (b) Density maps of $T$ = 4 shell hexamers from different CsoS2 truncation and I(V)TG mutation constructs. The map contour level is set to 4.5σ for all the maps except EMD-30385 (2σ).

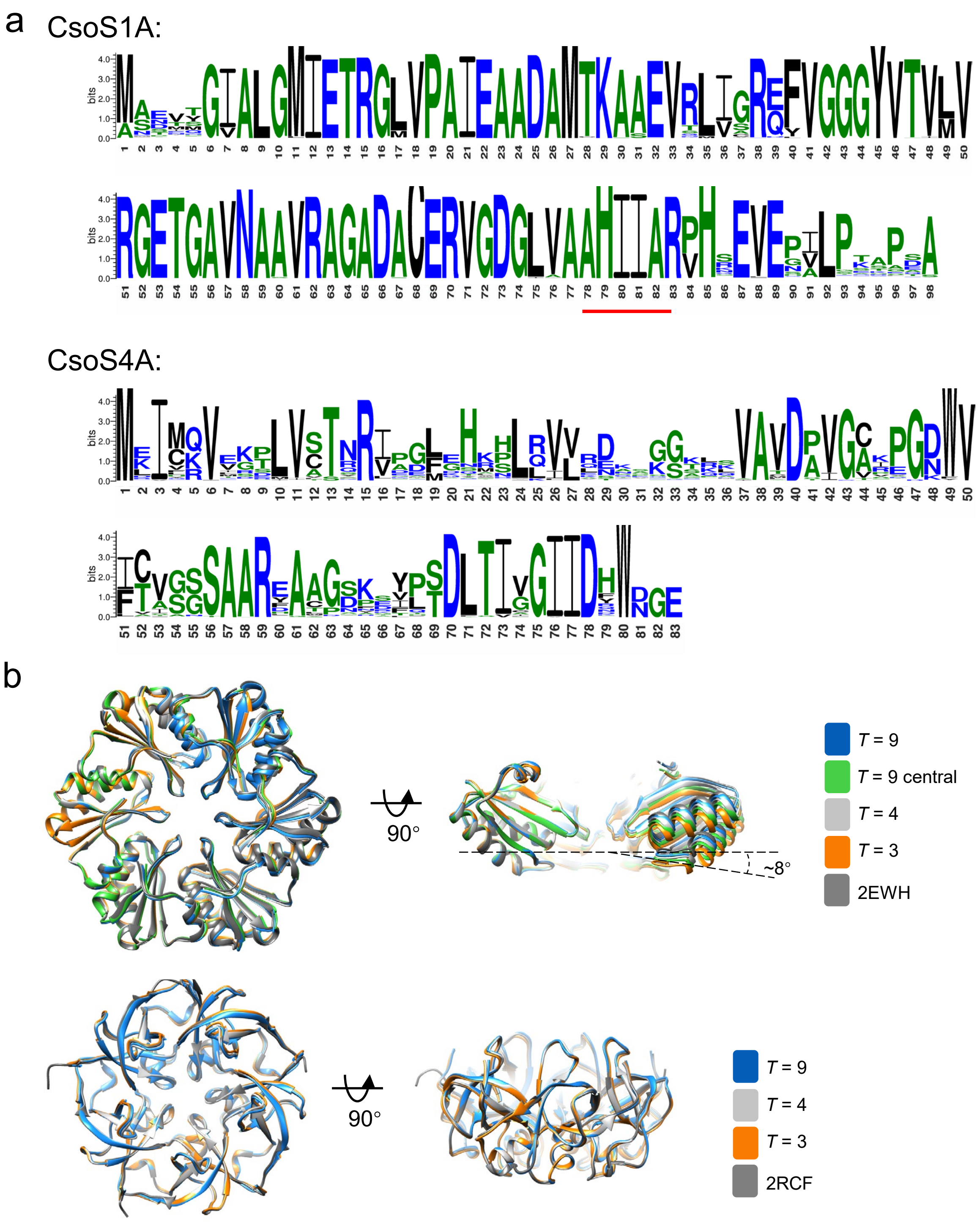


**Figure S5** | Sequence and structural analysis of CsoS1A and CsoS4A. (a) Conservation of CsoS1A (990 sequences) and CsoS4A (970 sequences) from the NCBI database, presented using Weblogo. The red line indicates the conserved β-strand interacting with the I(V)TG motif of CsoS2. (b) Structural comparison of CsoS1A hexamer (top) and CsoS4A pentamers (bottom) from mini-shell assemblies and X-ray crystallography structures in two orthogonal views, indicating very little deviations among these structures. Two quasi-equivalent hexamers from *T* = 9 shell are shown in blue (close to pentamer) and green (at the 3-fold), respectively (see Figure 2A). The hexamers in the *T* = 3 shell have the maximum curvature (~ 8°) compared with the crystal structure (PDB: 2EWH).

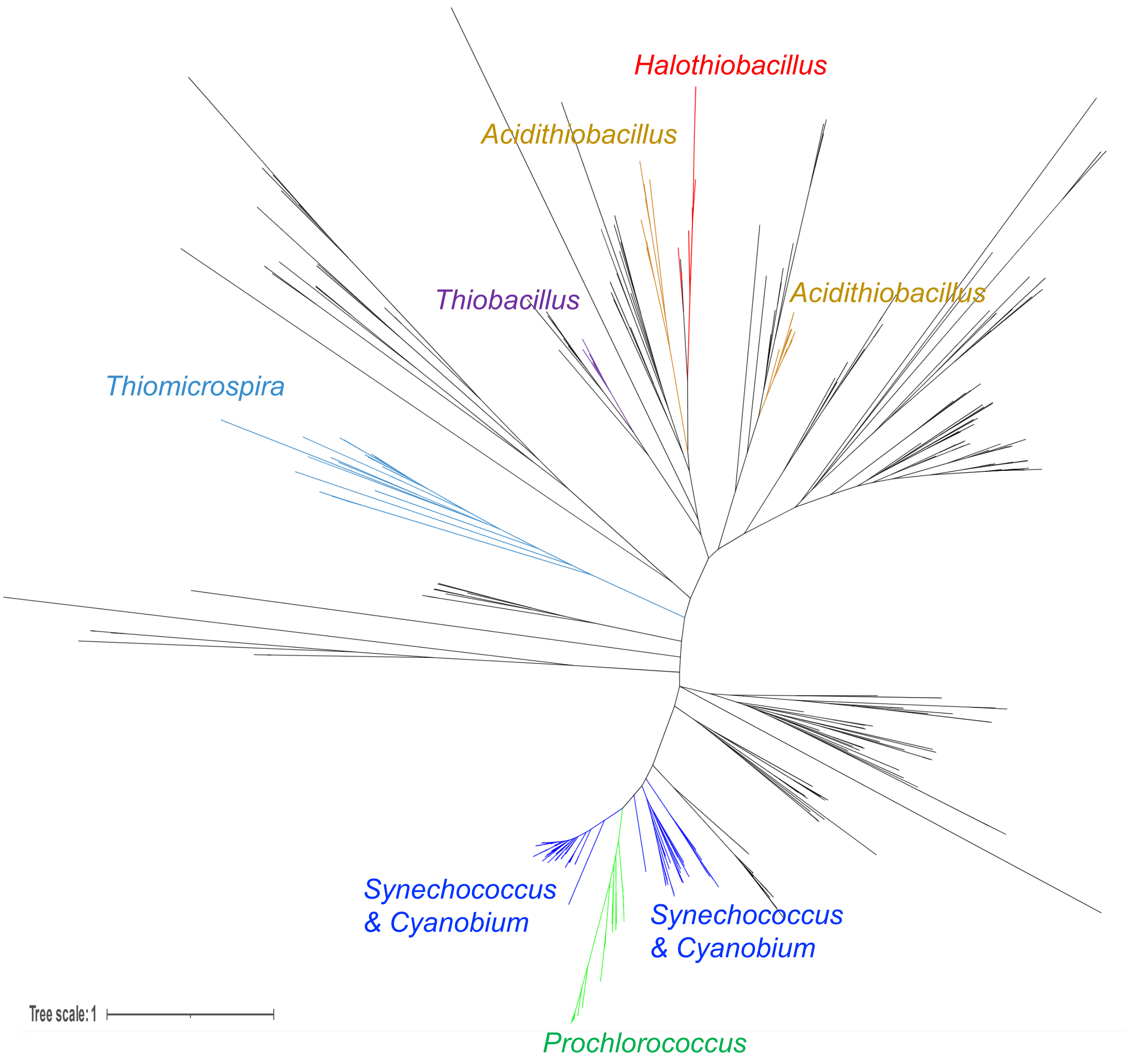


**Figure S6** | Maximum-likelihood phylogenetic tree of CsoS2. Among the 395 bacterial species containing CsoS2 homologs, clades of *Halothiobacillus, Thiobacillus, Thiomicrospira*, *Acidithiobacillus, Prochlorococcus, Synechococcus, and Cyanobium* are colored in red, violet, yellow, lollipop, green, and blue, respectively. Scale bar, 1 substitution per site.

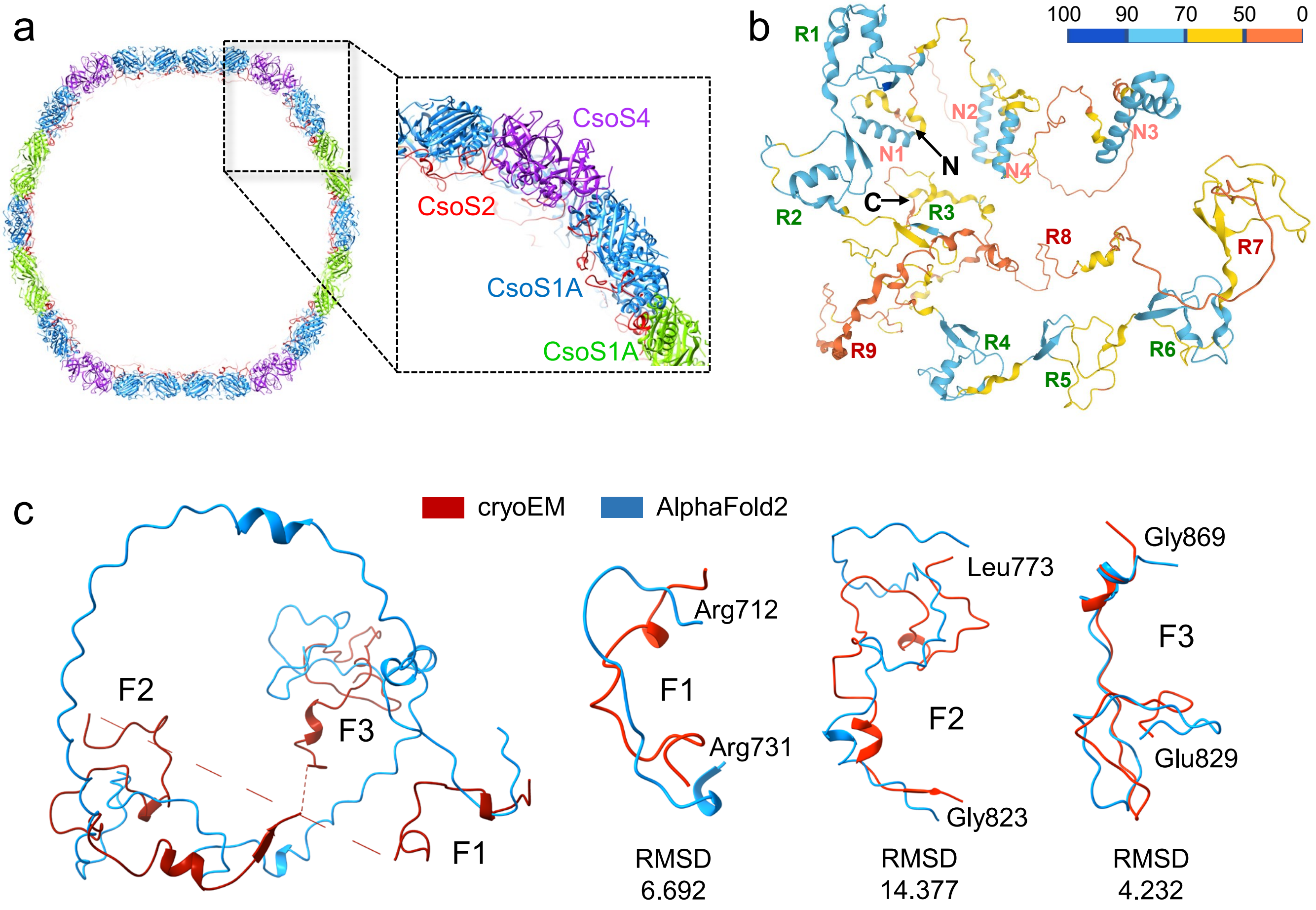


**Figure S7** | Identification and characterization of CsoS2 C-terminal fragments in mini-shell 2. (a) Localization of CsoS2 at the inner surface of $T$ = 9 shell assembly, shown as a central slice. Shell proteins are coloured the same as in Figure 1d. Inset shows a close-up view. (b) AlphaFold2 structure prediction of CsoS2. The predicted model is coloured according to model confidence scores (pLDDT) as indicated. The N- and C- termini and domains are labelled. (c) Overlay of CsoS2 structures from AlphaFold2 prediction (blue) and cryoEM (red). The structure of three individual fragments resolved by cryoEM and predicted from AlphaFold2 are overlaid on the right.

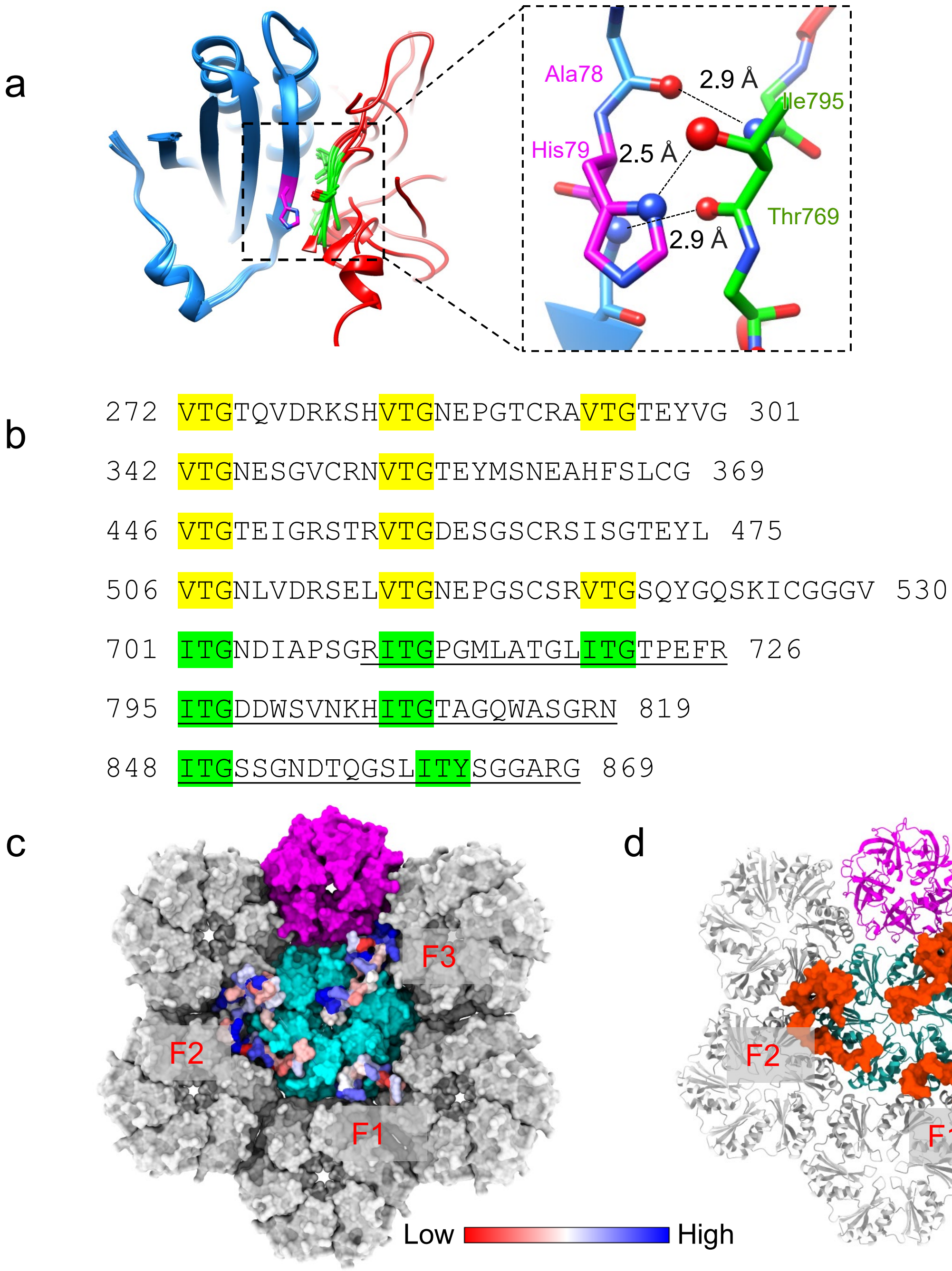


**Figure S8** | Conservation of CsoS2 C-terminal domain. (a) Hydrogen bond network between CsoS1A and a CsoS2 I(V)TG motif, mediated by both main-chain and side-chain hydrogen bonds. A closeup view of one of the [IV]TG motif interfaces (I795-T796-G797 in F2) is shown on the right. (b) [IV]TG motifs in *H. neapolitanus* CsoS2. The [IV]TG motifs in the Middle region and C-terminal domain are highlighted in yellow and green, respectively. The range of amino acid residues is labeled. (c) Surface rendering of the structure of CsoS2 C-terminal domain in complex with CsoS1A and CsoS4A in *T* = 9 shell. Only one copy of CsoS2 molecule is shown; the symmetry-related copies are removed for clarity. The CsoS1A hexamers are colored in cyan and gray, and the CsoS4A pentamer in purple. CsoS2 surface is colored according to conservation scores, ranging 1 to 10 (red to blue). The conservation score of CsoS2 fragments is calculated using ConSurf server (https://consurf.tau.ac.il/consurf_index.php). (d) Ribbon representation of the same structure shown in (b), with CsoS1A hexamers in gray/sea green, CsoS4A pentamer in purple and CsoS2 in red.

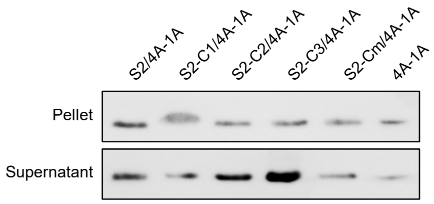


**Figure S9** | Western blot of mini shell constructs with different CsoS2 mutants. The assembled shell (in Pellet) and free shell proteins (in Supernatant) were probed with anti-CsoS1A antibody. The ratios of assembled shell and free shell proteins are examined to indicate shell assembly efficiency (see Fig. 4b).

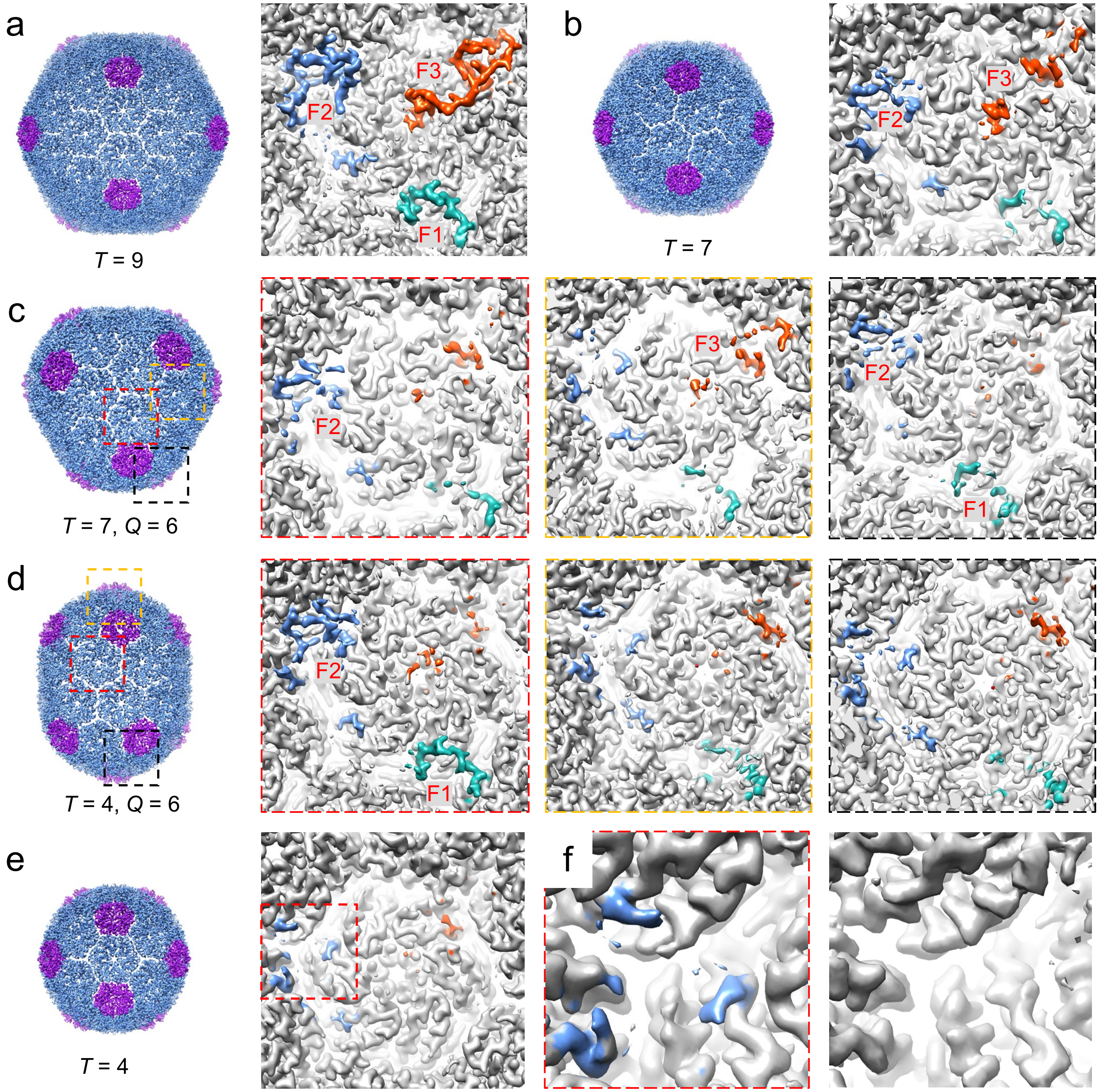


**Figure S10** | CsoS2 density in shell assemblies from the mini-shell 4 construct containing S2-C1). (a) *T* = 9 shell with a close-up view of CsoS2 F1, F2 and F3 fragments, coloured in light sea green, blue and orange, respectively. (b) *T* = 7 shell with a close-up view of CsoS2 F1, F2 and F3 fragments, where only F2 and F3 fragments can be assigned confidently, albeit with weaker density. (c) *T* = 7, *Q* = 6 shell with close-up views of three quasi-equivalent interfaces. The densities corresponding to [IV]TG motifs can be observed. (d) *T* = 4, *Q* = 6 shell with close-up views of three quasi-equivalent interfaces. CsoS2 densities are coloured according to their respective fragments. (e) *T* = 4 shell with a close-up view. (f) Comparison of *T* = 4 shells from mini-shell 2 (left) and mini-shell 1 (right). The densities of CsoS2 are colored in blue. Only the residual densities corresponding to [IV]TG motifs can be observed.

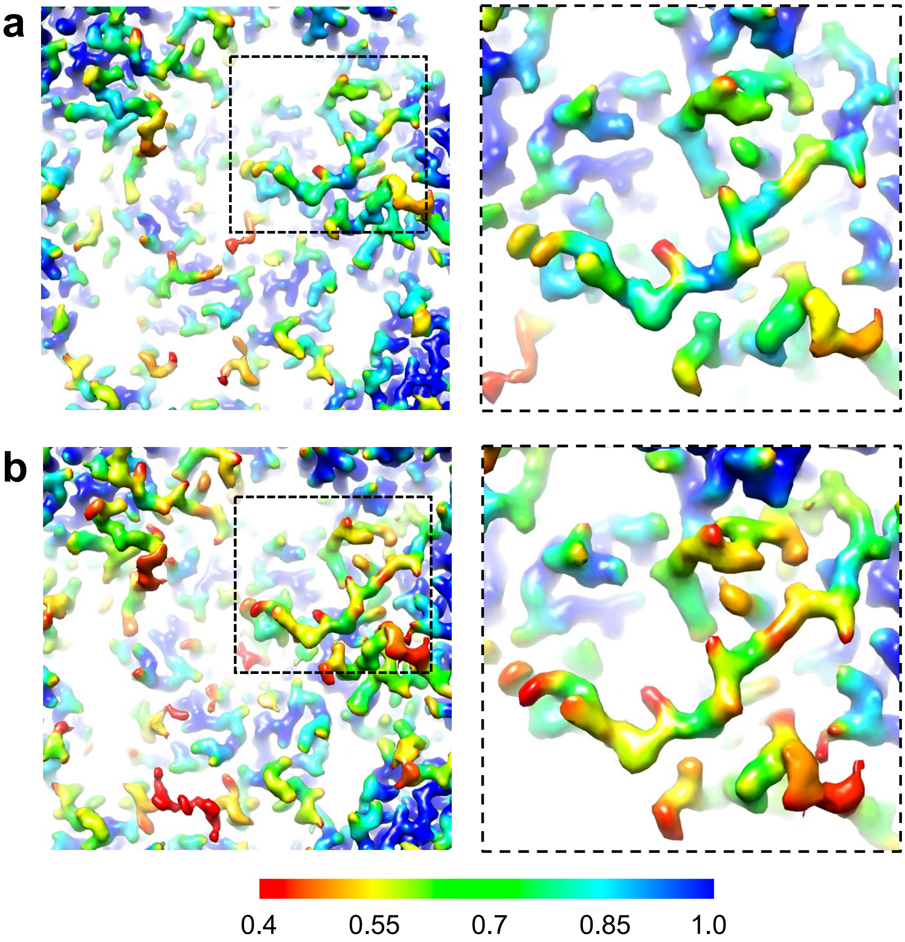


**Figure S11** | Quantification of CsoS2 occupancy by Occupy. (a) Occupancy map of T=9 shell from the full-length CsoS2 construct, with a F3 fragment zoomed in (right). (b) Occupancy map of T=9 shell from the truncated C1 construct, with a F3 fragment zoomed in (right). Density map of a hexamer from the T=9 shell colored by occupancy from 0.4 to 1 (red to green). F3 fragment in T=9 C1 construct has lower occupancy.
